# Stochastic population growth and its implications for mutation dynamics under carrying capacity constraints

**DOI:** 10.64898/2026.10.21.760709

**Authors:** Jon Bohlin, John H.-O. Pettersson

## Abstract

Mathematical models of mutation processes often assume deterministic population growth, although biological populations are continuously subjected to random fluctuations. Here, we extend a model inspired by the classical Luria–Delbrück framework by introducing stochastic perturbations into population growth while assuming a constant mutation rate. Using Itô calculus, we derive an analytical solution for a stochastic logistic growth model with carrying capacity. By comparing the stochastic model with its deterministic counterpart, we demonstrate that random perturbations can substantially influence population variability, particularly as the population approaches its carrying capacity. While deterministic growth levels off, stochastic fluctuations persist and may become increasingly important relative to the underlying growth dynamics. The magnitude of these effects depends on the relationship between population growth and stochastic volatility. We illustrate the model using real life within-host SARS-CoV-2 infection data and discuss its implications. Our findings highlight the importance of accounting for stochastic population dynamics when modelling mutation-related processes in capacity-limited microbial populations.

## Introduction

The classical Luria–Delbrück fluctuation experiment [21] demonstrated that mutations conferring bacteriophage resistance could arise spontaneously before exposure to selective pressure. The experiment subsequently provided the foundation for mathematical models describing mutation processes in growing microbial populations. Although numerous extensions have accounted for different aspects of mutation and population dynamics, assumptions concerning population growth remain important for the interpretation of these models. The intuition behind the Luria-Delbrück model, and in particular the subsequent extension of it by Lea and Coulson [18], was that there is a probability that a bacterium will mutate during replication [21]. This was a reasonable assumption as mutations often occur during replication [12]. However, mutations also occur in-between cell divisions [3]. In particular, methylation and subsequent deamination of Cytosine (C) resulting in Thymine (T) is largely responsible for the over-representation of A + T mutations as compared to Guanine (G) and C mutations that are more often fixed [3, 11]. Hence, mutation rates are sometimes better modeled with respect to time instead of per generation [33].

Several updates have been added to the original Luria-Delbrück model [6, 27, 16, 13, 31, 32, 33, 34] and some quirks have been found and corrected [30]. Although the Luria–Delbrück experiment provided compelling statistical evidence that phage resistance can arise before exposure to selective pressure, the molecular mechanisms underlying bacteriophage resistance are considerably more complex than those represented by classical mutation models. Indeed, recent work has demonstrated that bacteriophage T1, historically considered strictly lytic, can establish a lysogenic state under certain conditions, with its transition to lysis influenced by the metabolic and signalling state of its bacterial host [17, 7, 14]. These findings do not invalidate the classical interpretation of the fluctuation experiment but emphasize that the biological mechanisms underlying resistant phenotypes may be more diverse than those assumed in mathematical models. Nevertheless, the original model, together with the added refinements, gives a good intuition of mutation processes in non-recombining organisms thanks to its simplicity and intuitive reasoning.

In the present study, we investigate how multiplicative stochastic perturbations in population growth influence mutation-related quantities under carrying-capacity constraints. Using an Itô stochastic differential equation [35], we derive the population dynamics and examine their moments, with particular emphasis on the relationship between deterministic growth and stochastic volatility. We then compare these results with the corresponding deterministic model and discuss their potential implications for within-host microbial and viral evolution. We follow the reasoning of Zheng [33] and the original Luria-Delbrück [21] article in motivating our models. Demonstration of the models are provided on a class of growth models with populations subjected to a carrying capacity, something that is more realistic when modeling microbial, including viral, populations and compare with the corresponding deterministic counterparts. We also provide a practical example based on within-host mutations of the SARS-CoV-2 virus [29].

## Mathematical models

### The classical Luria-Delbrück model

The classical Luria-Delbrück model [21, 33] states that the number of mutations in a plated culture can be formulated as the mutation-event rate per individual per unit time *µ, i*.*e*. the fraction representing mutations with respect to generation or time *t*, multiplied by a growth equation of population *N* :

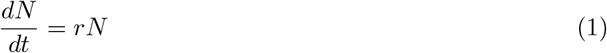

where *r* is the growth rate and *N*_0_ is assumed to be the initial population size. In the following, we assume a constant mutation-event rate *µ* per population member per unit time. The quantity *m*(*t*) = *µN* (*t*) therefore represents the instantaneous expected rate of mutation events. These quantities do not explicitly account for the subsequent reproduction, selection, or loss of mutant lineages. Our primary interest is how stochastic population growth influences these mutation-related quantities. As the solution to this equation is *N* (*t*) = *N*_0_ exp (*rt*), the instantaneous expected rate of mutation events in the population at generation or time *t* can thus be written as:

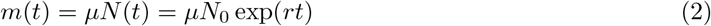

Accumulated mutations is of interest to understand how natural selection operates on a population at the molecular level over time. Experiments such as the long time evolution experiment (LTEE) [5] has demonstrated this. The cumulative expected number of mutation events up to generation or time *t*, conditional on the specified population trajectory, can be formulated as follows:

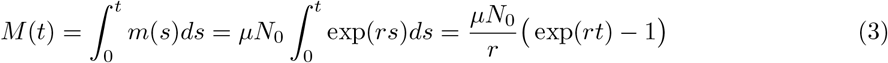

The equations above are similar to how the original Luria-Delbrück model was formulated [21]. However, it is unrealistic to model a bacterial population as indefinitely increasing, especially when focusing on infections within a host [25]. Indeed, models concerned with bacterial growth are sometimes divided into three phases [1]: lag phase, exponential phase and stationary phase. We shall therefore in the following assume that a population grows until it reaches a carrying capacity *K* which is taken here to be the stationary phase. In the following we will also exclusively focus on mutation rates with respect to time and not generation.

### The Luria-Delbrück model based on a constrained population size

If we assume that the population modeled *N* has a carrying capacity *K* and a growth rate *r* with respect to time *t* [4, 1] we get:

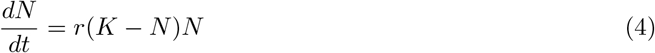

It should be noted that, 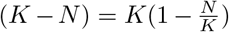. While eq.(4) is a non-linear differential equation it can still be solved analytically (See Appendix) giving:

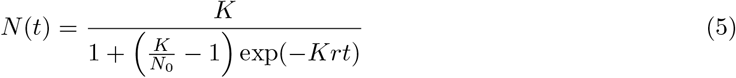

The number of mutations in a population with carrying capacity *K* at time *t* will now be stated as

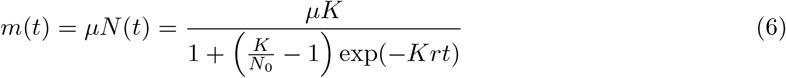

and the corresponding formula for accumulated mutations *M* (*t*) is then:

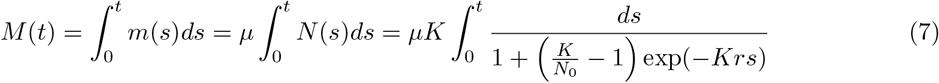

after a bit of straight-forward algebra (see Appendix) we get a closed-form solution for the number of accumulated mutations over time *t*:

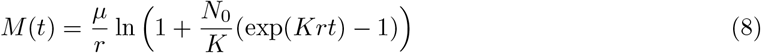

for positive *r, K* and *N*_0_.

### A Luria-Delbrück model based on Itô calculus

The simple growth model described by eq.(1) assumed a fixed growth rate *r*. Populations of living organisms tend to vary in size and so allowing growth to be influenced by stochastic events would make the model more realistic. If we instead let the growth rate be *r* = *κ* + *cW*_*t*_(*ω*), where *κ* is the deterministic growth rate, *W*_*t*_ is a Gaussian white noise with respect to time *t* and random trajectory *ω ∈* Ω, multiplied by the scaling coefficient *c*, and put this into eq.(1), we get:

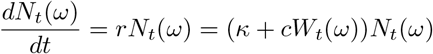

*N*_*t*_(*ω*) is now the population with respect to time *t* and trajectory *ω*. It can be shown [35] that the simple stochastic growth model can be formulated as a stochastic differential equation (SDE):

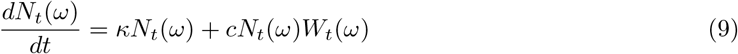

The SDE can be solved using methods from stochastic integration and in the present work we will solely employ Itô calculus. Eq.(9) can now be written on differential form with respect to time *t*:

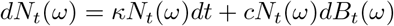

*B*_*t*_(*ω*) represents Brownian motion and 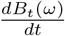 is interpreted as white noise *W*_*t*_(*ω*). Where it is clear that the equations discussed are stochastic we shall assume the variable *ω* implicitly. The solution to this SDE is straight-forward and has been elaborated in detail elsewhere [35] but we also give a sketch of its solution in the Appendix:

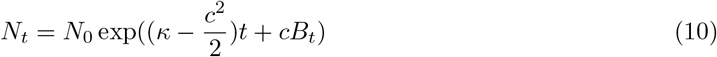

*N*_*t*_ is now assumed to be a stochastic process adapted to a filtration *ℱ*_*t*_ which in turn consists of increasing *σ*-algebras of sets *A ∈* Ω up to time *t*. In other words, the filtration *ℱ*_*t*_ contains the history of a population up to time *t. N*_0_ is a random variable independent of Brownian motion.

The conditional mutation-event intensity, and thus the stochastic version of the Luria-Delbrück model, can then be formulated analogously to eq.(2):

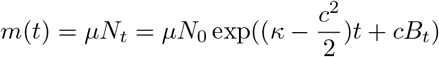

and similarly to eq.(3) for the time-integrated intensity:

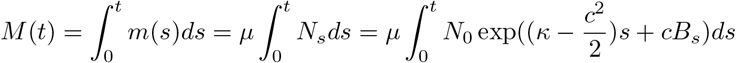

The integrals and corresponding measures *ds* are considered to be of Lebesgue type with respect to Borel *σ*-algebras in ℝ^+^.

### An integrating-factor approach for the solution of a class of nonlinear SDEs

Quite remarkably, there exist different formulas for solving classes of SDEs in spite of an additional Brownian motion term. Moreover, since stochastic integration does not conform to the assumption of bounded variation (but to bounded quadratic variation), the Brownian motion term will vary according to the stochastic integration method used (*i*.*e*. Itô or Stratonovich) [26]. Here we will primarily focus on the Itô integral which can be transformed to the Stratonovich integral (and vice versa)[26]. In the following we will assume that all stochastic processes are adapted to some filtration ℱ_*t*_ consisting of *σ*-algebras of sets *A* Ω. All non-stochastic integrals are assumed to be of Lebesgue type with respect to *σ*-algebras of sets *U ∈* ℝ^+^.

To state a SDE version of the growth equation with a carrying capacity *K* (eq.(4)) we will need some smaller results [35]. We write out the different results explicitly here.

First, we motivate a formula for partial integration of Itô processes *X*_*t*_ and *Y*_*t*_. We will in general avoid the use of cross-variation processes (see [26, 15]) and follow the more simplified and accessible notation used by Øksendal [35] for clarity. By using the multi-dimensional Itô formula [35, 26] on 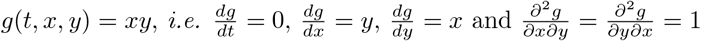 so that:

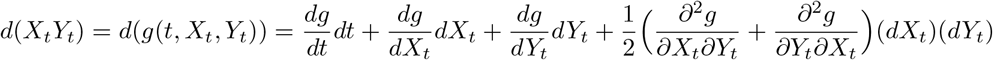

we get a formula for partial integration of Itô processes:

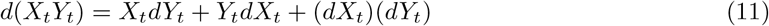

The exponential martingale:

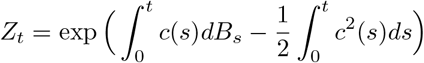

is important in stochastic analysis not only because it is a martingale but also due to its many applications and theoretical significance (see [35]). We are interested in its inverse:

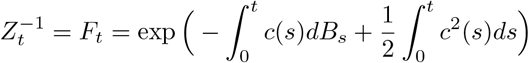

and its differential which can be found by setting *g*(*t, x*) = *e*^*x*^ and:

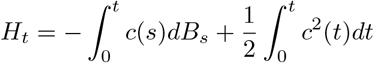

Thus,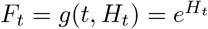. Using Itô’s formula we get:

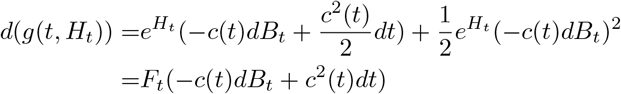

so that:

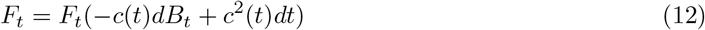

If we now use eq.(11) on *d*(*F*_*t*_*X*_*t*_) with *dX*_*t*_ = *f* (*t, X*_*t*_)*dt* + *c*(*t*)*X*_*t*_*dB*_*t*_ we have:

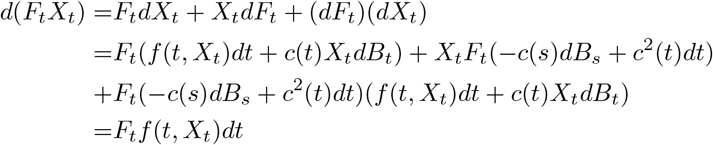

By setting:

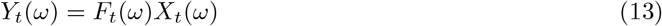

for each trajectory *ω ∈* Ω the general formula is:

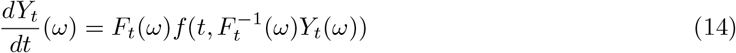

for solving SDEs on the form:

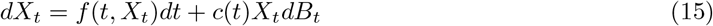

where *f* : ℝ_+_ *×* ℝ *→* ℝ and *c* : ℝ *→* ℝ are assumed to be continuous while

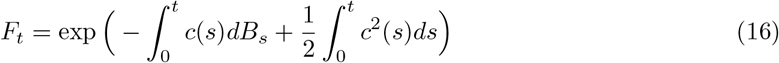

is the integrating factor.

### Stochastic growth of a population with a carrying capacity

Now that we have a general formula for solving a class of non-linear SDEs it can be applied to the stochastic analog of population growth with carrying capacity *K* (eq.(4)), *i*.*e*.

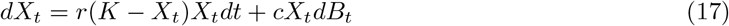

Let 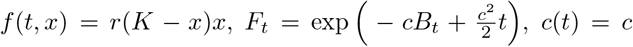 is assumed to be a constant, and set *Y*_*t*_ = *F*_*t*_*X*_*t*_. If this is put into eq.(14) we have:

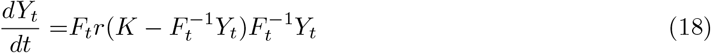

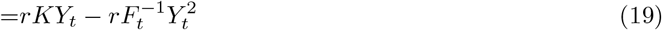

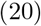

Since 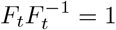 Let 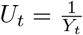. As eq.(18) does not have a *dB*_*t*_ term:

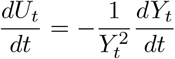

Substituting for 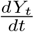:

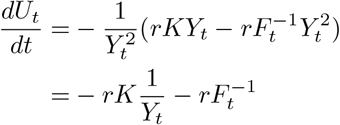

or, equivalently,

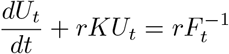

Since this is a standard linear SDE it can be solved using the integrating factor *e*^*rKt*^ so that

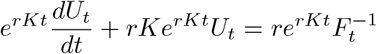

which is

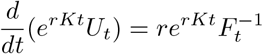

Integrating from 0 to *t* and rearranging implies:

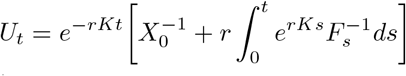

where we have used that 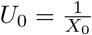. Since *U*_*t*_ = 1*/Y*_*t*_, substitution gives:

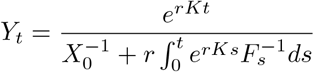

We have that 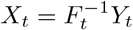, therefore:

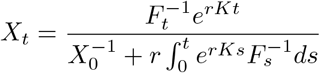

Since 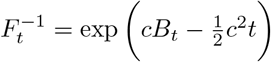, the numerator becomes:

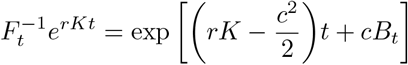

and so:

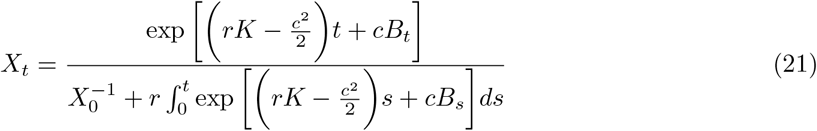

In form, eq.(21) is similar to eq.(5) above. At the cost of considerably more calculations it is possible to solve eq.(17) differently without using formula (14). Although more elaborate, the steps are straightforward given the following procedure; using the Itô formula with the integrating function 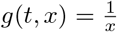 gives a linear SDE with respect to 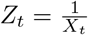. This linear SDE can then be solved by using the Itô formula again together with eq.(11) on *Z*_t_ and integrating factor 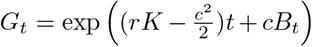.

The deterministic analog to eq.(21) is:

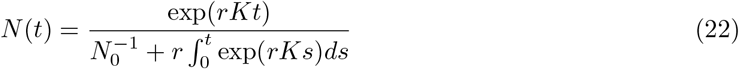

and can be found by using the chain-rule on 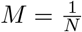 and the fact that 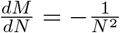 in effect linearizing eq.(4) analogously to eq.(17) and the Itô formula as described above. From these solutions it can be seen that the stochastic model differs from the deterministic via the subtraction of 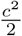, in addition to the volatility term *cB*_*t*_. When comparing the stochastic-with the deterministic Luria-Delbrück model, with a population having a carrying capacity *K*, we will focus on eq.(21), and its deterministic counterpart eq.(22), for ease of interpretation. The stochastic Luria-Delbrück model, and its deterministic counterpart, can be seen in Figure 1.

**Figure 1.**
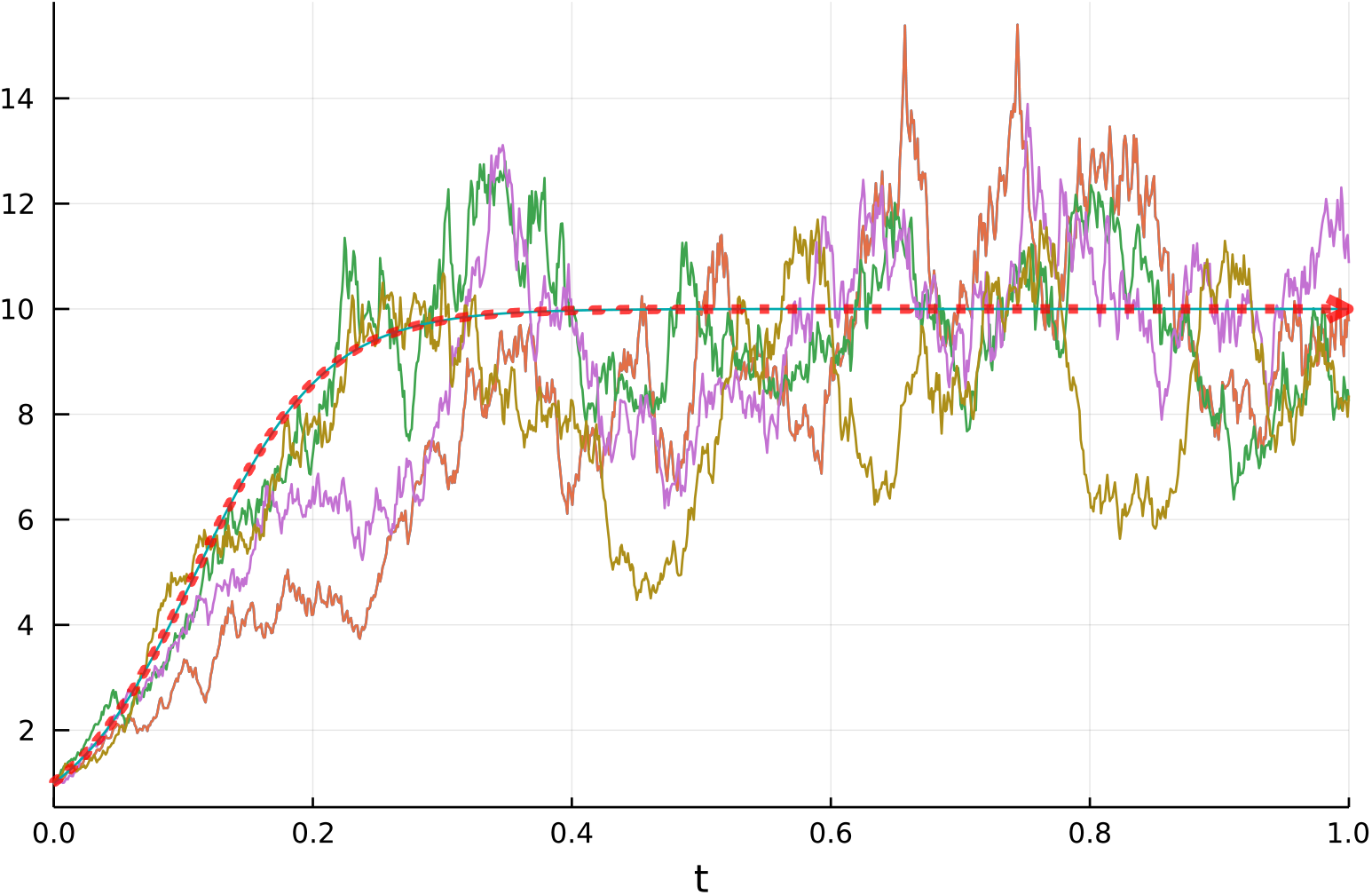
Five realizations of the Luria-Delbrûck model based on a stochastic population with a carrying capacity *K* with the parameters *K* = 10, 0 *≤t ≤*1, *c* = 1.0, *r* = 2.0 and *X*_0_ = 1 together with the deterministic counterpart (red dotted line). The vertical axis designates the number of mutations *m*(*t*) while the horizontal axis represents time *t*.

### Expectation and Variance of *X*_*t*_

While a closed form expression of the expectation of *X*_*t*_, as described by equation (21), is impossible some simplifications can be made. Let 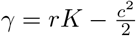. This gives:

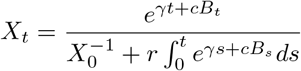

the numerator can be rewritten as:

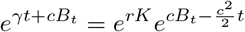

This is a term with the exponential martingale 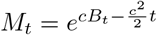 so that 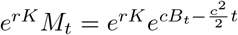 Eq.(21) can then be written as the expectation:

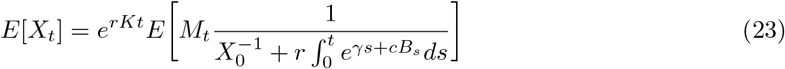

Due to the martingale *M*_*t*_ the Girsanov transform can be employed with the Brownian motion:

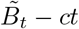

This can now be inserted into the eq.(23) so that

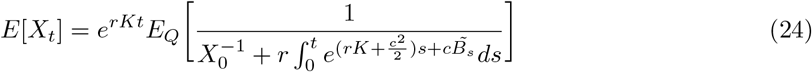

due to the change-of-measures formula. The variance can now be calculated using:

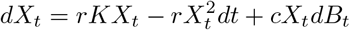

and that

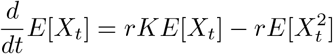

Let *m*(*t*) = *E*[*X*_*t*_] so that 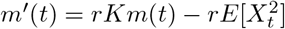 implying that:

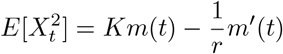

The variance 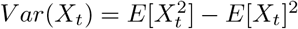 can now be stated as:

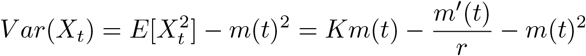

This relation can be stated more explicitly by using eq.(24):

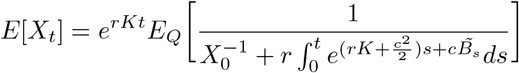

and the fact that

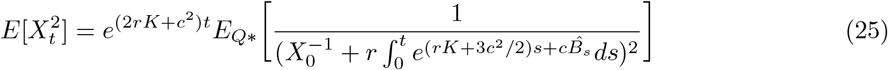

which is due to 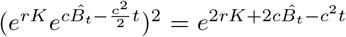 and particularly that 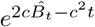 also defines a change of measure which ultimately gives (25) and hence:

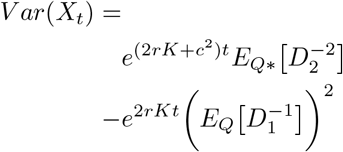

where

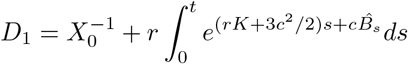

and

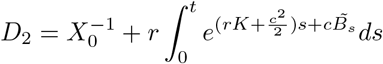

### Simulation and presentations of models

The ODEs and SDEs seen in the figures below were computed with Julia version 1.6.1 [2] using the “DifferentialEquations” library. The fifth order adaptive time stepping method (“Tsit5”) was used as the solver for ODEs while the Euler-Maruyama (“EM”) solver was used for SDEs. All figures were also made with Julia and the “Plots” library.

## Results and Discussion

### Implications of growth and carrying capacity in a stochastic population

It can be seen from eq.(21) that the stochastic effects can have profound influence on growth particularly if growth rates are low. That is, when 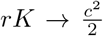. The stochastic effects will then influence growth in a feedback-like manner affecting both growth- and population size decline as can be seen in the illustrative simulations in Figure 2.

**Figure 2.**
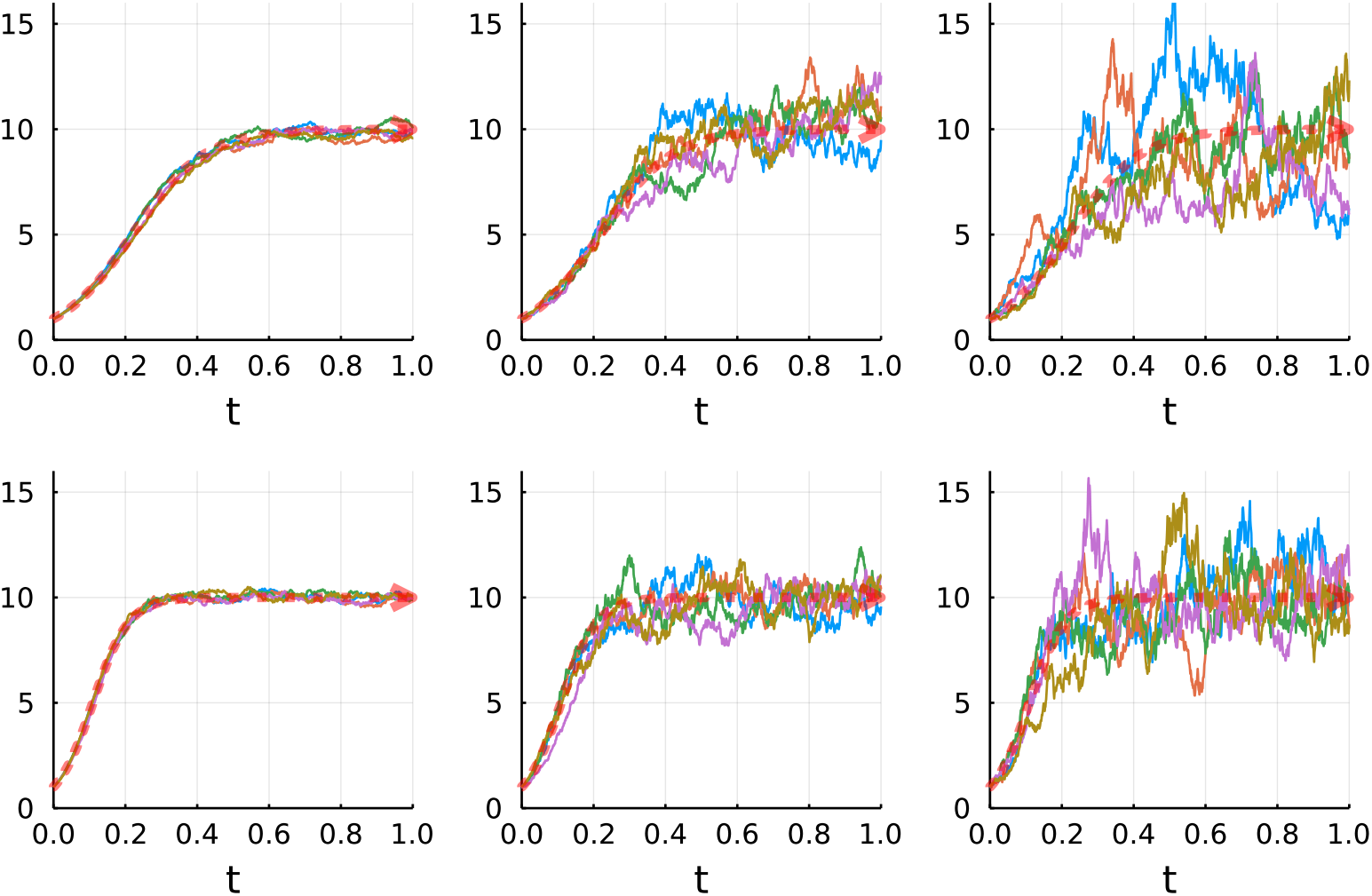
Five realizations of the stochastic Luria-Delbrûck model based on a population with carrying capacity *K* = 10 and parameters set to *β* = *{* 0.1, 0.5, 1 *}* (from left to right), *r* = *{*1, 2 *}* (top to bottom), *µ* = 1 and *X*_0_ = 1. The vertical axis designates the number of mutations *m*(*t*) while the horizontal axis represents time 0 *≤t ≤*1. The red dotted line represents the deterministic model.

As the population approaches its carrying capacity, the deterministic growth term gradually decreases, whereas the stochastic perturbations remain active. Consequently, random fluctuations become increasingly important relative to the deterministic growth dynamics. Unlike the deterministic population, which approaches a stable equilibrium, the stochastic population may continue to fluctuate around the carrying-capacity level. The magnitude of these fluctuations depends on the relationship between the deterministic growth parameters and the volatility of the stochastic process (see Figure 2).

For positive *X*_0_, *r, K, c*, assuming:

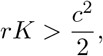

the stochastic logistic model eq.(21) has a nondegenerate stationary distribution [20]. It is a gamma distribution, with stationary mean

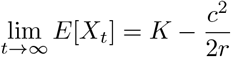

and stationary variance

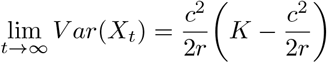

where lim_*t→∞*_ *X*_*t*_ denotes a random variable *X*_*∞*_ with the limiting stationary distribution. This follows from the stationary Fokker-Planck equation and is consistent with known results for logistic growth under multiplicative environmental noise [20]. It also shows that the stochastic population described by eq.(21) has a stationary mean below *K*, even though the deterministic model approaches *K* itself. For *rK ≤ c*^2^*/*2, it is clear that eq.(21) instead approaches extinction in the long-time limit, under the usual positive-initial-population assumptions.

### An example involving SARS-CoV-2

The virus responsible for the SARS-CoV-2 pandemic has a positive sense single stranded RNA genome. It belongs to the sub-genus Sarbecovirus of the Betacoronavirus genus and the size of its genome is approximately 30,000 nucleotides [29]. From its identification late 2019, the virus has evolved into several variants of concern (VoC) [10]. Numerous studies have investigated how these VoCs have evolved (see [10] and reference therein) but there are still many unanswered questions regarding how the virus evolves within a host [22, 28, 8]. The mechanisms responsible for how variants are transferred between hosts is also not completely understood. Indeed, the SARS-CoV-2 genomes isolated during infection are in reality collapsed consensus sequences stemming from numerous different variants [19]. Since mutation rates estimated for the SARS-CoV-2 virus [24] are based on consensus sequences transmitted between infected they may not be reflective of actual within-host mutation rates. Indeed, by setting parameters reflective of how SARS-CoV-2 grows within a host [23] in the deterministic and stochastic Luria-Delbrück models we find that the number of mutations are likely somewhat high (see Figure 3) compared to most published studies [28, 22, 9]. Of course, the number of within-host variants depend on several parameters. For instance, many within-host SARS-CoV-2 variants may be selected against during the bottleneck event [28] and thus before host-to-host transmission. Others still, may carry strongly fitness decreasing mutations that are selected against after several replications but before the bottleneck event. Moreover, assuming that a representative sample of within-host SARS-CoV-2 diversity can be obtained, the sampled diversity will likely also depend on timing (see figures 1 - 3). Indeed, variants collected after the immune system is activated are likely subjected to very different selective pressures compared to those variants present prior to immune activation [25]. Selective pressures will likely also impact growth and subject the viral population to stochastic effects that in turn will reflect viral diversity [23, 25]. As was seen above, the Luria-Delbrück formula states that if the growth rate is high compared to the stochastic volatility, genetic drift will only exert a small influence during viral growth. A reduction of growth, say through the action of the hosts’ immune defence system, will increase the influence of the stochastic volatility term that could potentially have an effect on viral diversity as has recently been reported [25, 9]. From the model presented here (eq.(21)), the stochastic effects are mostly affecting diversity after the viral population has reached its carrying capacity. The selective constrains responsible for the carrying capacity of SARS-CoV-2 within a host likely vary between hosts and even cell types [25]. Furthermore, there are indications that an impaired immune reaction to the virus decrease the number of virions but increase genetic diversity [28, 22]. Hence, including a vanishing volatility term in the mutation model introduces a set of assumptions and should therefore be properly justified.

**Figure 3.**
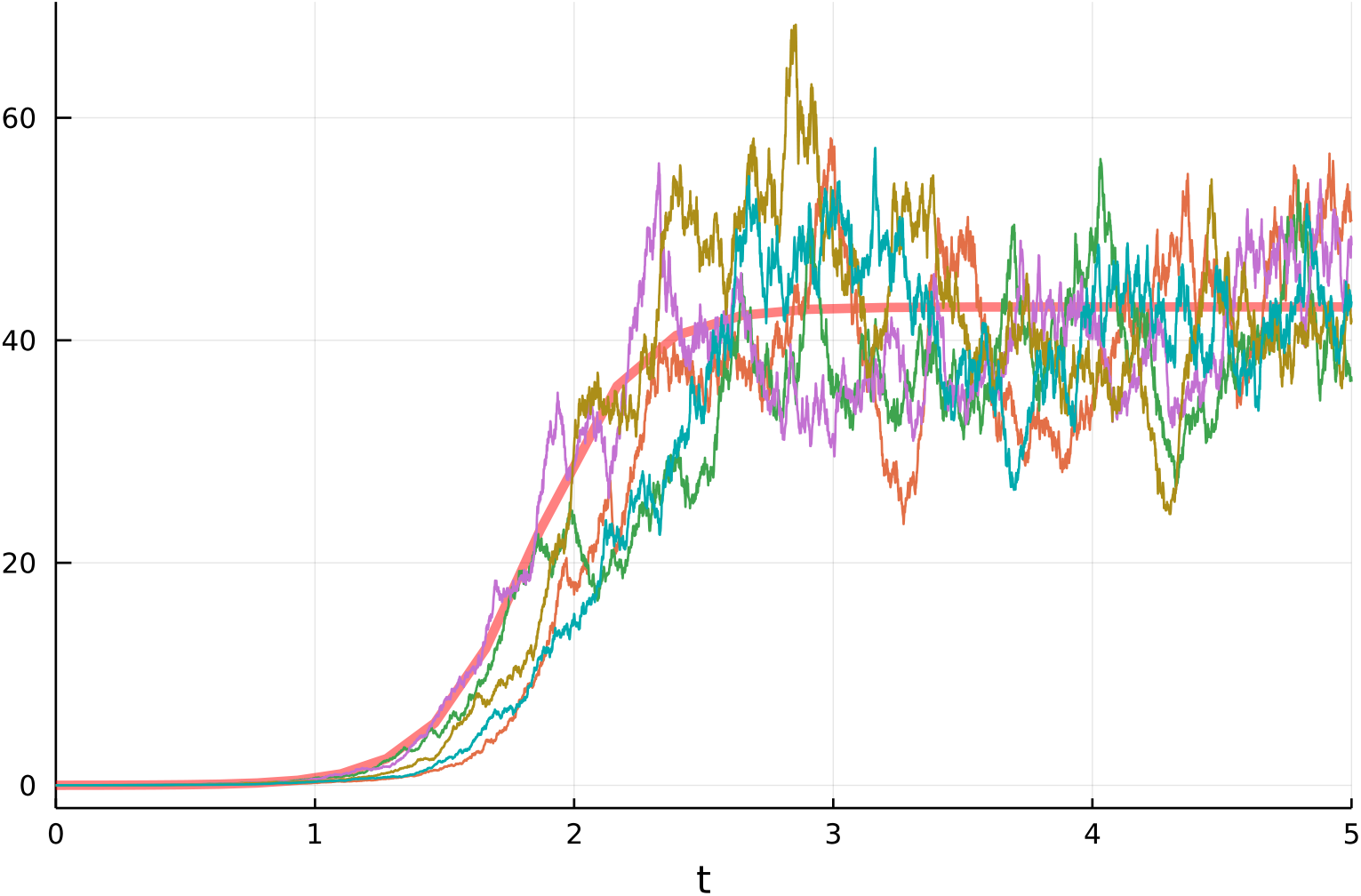
Simulated mutation-event intensity in a within-host SARS-CoV-2 population under deterministic and stochastic logistic growth. Carrying capacity is assumed to be *K* = 10^7^ virions, with starting concentration *X*_0_ = 1000, growth rate *r* = 0.5 *·* 10^*−*6^ and mutation rate *µ* = 4.38 *·*10^*−*6^. Time *t* is in days. The thick red line designates the deterministic growth equation, the differently colored lines are stochastic realizations. Volatility is set to *β* = 0.5 for the stochastic model.

The SARS-CoV-2 example is intended to illustrate the potential relevance of stochastic population growth to within-host viral evolution, rather than to provide a calibrated model of within-host genetic diversity. In particular, the model does not explicitly incorporate selection, transmission bottlenecks, or differences in replication rates between viral variants. Consequently, simulated mutation-event intensities should not be interpreted directly as the number of distinct variants detectable through viral genome sequencing.

## Conclusion

Here we present an Itô calculus based version of the classical Luria-Delbrück mutation model. We demonstrate this model for a population with simple stochastic growth. Furthermore, we provide a simple formula for a general class of non-linear stochastic differential equations and use it to demonstrate the model on a stochastic population with a carrying capacity. The stochastic models are compared with their corresponding deterministic counter parts. We find that the Itô calculus based Luria-Delbrück model incorporates stochastic events in a natural way. Moreover, the models presented here demonstrate that if the population growth-rate decreases below a given threshold the stochastic processes may dominate and in turn influence genome evolution in a feedback-like manner. For the model incorporating a stochastic population with a carrying capacity it is demonstrated that the stochastic effects increase as the population reaches its carrying capacity. We end by discussing these findings in the light of the SARS-CoV-2 virus.

## Declarations

## Ethics approval and consent to participate

Not applicable

## Consent for publication

Not applicable

## Availability of data and materials

Not applicable

## Authors’ contributions

Conception: JB & JHOP Mathematical model: JB Evolutionary analysis: JB & JHOP Wrote the manuscript: JB

Both authors approved the final manuscript.

## Acknowledgements

ChatGPT Version 5.6 Sol is credited for the section “Expectation and Variance of *X*_*t*_ and for suggesting Eq.(18) and the authors take full responsibility for these results.

