## Appendix for "Stochastic population growth and its implications for mutation dynamics under carrying capacity constraints"

### Appendix: Exploring the influence of mutation rates with stochastic perturbations on genome evolution

October 30, 2026

#### The Luria-Delbrück model based on a constrained population size

Solving the deterministic growth model with carrying capacity K:

$$\frac{dN}{dt} = r(K - N)Ndt \tag{1}$$

Note the relation:

$$\frac{1}{N} + \frac{1}{K - N} = \frac{K - N}{(K - N)N} + \frac{N}{(K - N)N} = \frac{K}{(K - N)N}$$

Allowing (1) to be written as:

$$\Big(\frac{1}{N} + \frac{1}{K - N}\Big)dN = Krdt$$

The integral can now be computed on both sides using the standard chain rule:

$$\ln\Big(\frac{N}{K - N}\Big)\Big|_0^t = Krt$$

After some straight-forward algebra we have the final formula:

$$N(t) = \frac{K}{1 + \Big(\frac{K}{N_0} - 1\Big)\exp(-Krt)} \tag{2}$$

##### The number of accumulated mutations

To see that the closed form formula for the integral representing the number of accumulated mutations over time t is:

$$M(t) = \int_0^t m(s)ds = \mu\int_0^t N(s)ds = \mu K\int_0^t \frac{ds}{1 + \Big(\frac{K}{N_0} - 1\Big)\exp(-Krs)} = -\frac{\mu}{r}\ln\Big(\frac{1}{1 - \frac{N_0}{K}(1 + \exp(Krt))}\Big) \tag{3}$$

Let $u = \Big(\frac{K}{N_0} - 1\Big)\exp(-Krs)$, and use the fact that:

$$\frac{1}{(1 + u)u} = \frac{1}{u} - \frac{1}{1 + u}$$

The chain rule can then be applied with respect to $\frac{du}{ds}$. Proceeding analogously as above gives the closed formula:

$$\mu K\int \frac{ds}{1 + \Big(\frac{K}{N_0} - 1\Big)\exp(-Krs)} = -\frac{\mu}{r}\int \frac{du}{u(1 + u)} = -\frac{\mu}{r}\int \Big(\frac{1}{u} - \frac{1}{1 + u}\Big)du$$

and thus the anti-derivative with respect to $u$ is

$$-\frac{\mu}{r}\ln\Big(\frac{u}{1+u}\Big)$$

Inserting for $u$, taking limits and re-arranging:

$$-\frac{\mu}{r}\ln\Big(\frac{\Big(\frac{K}{N_0}-1\Big)\exp(-Krs)}{1+\Big(\frac{K}{N_0}-1\Big)\exp(-Krs)}\Big)\Big|_0^t$$

gives the desired result:

$$\frac{\mu}{r}\ln\Big(1+\frac{N_0}{K}(\exp(Krt)-1)\Big)$$

### A Luria-Delbrück model based on Itô calculus

Solution to the simple stochastic growth model:

$$dN_t(\omega)=\kappa N_t(\omega)+cN_t(\omega)dB_t(\omega)$$

It is the clear that:

$$\frac{dN_t}{N_t}=\kappa dt+cdB_t$$

and thus

$$\int_0^t \frac{dN_s}{N_s}=\kappa t+cB_t$$

The Itô formula can now be used for $g(t,x)=\ln x$

So we get:

$$d(\ln N_t)=\frac{1}{N_t}dN_t+\frac{1}{2}\Big(-\frac{1}{N_t^2}\Big)(dN_t)^2$$

Since

$$dN_t=\kappa N_t+cN_tdB_t$$

inserting this into the equation above we get

$$d(\ln N_t)=\frac{dN_t}{N_t}-\frac{1}{2N_t^2}c^2N_t^2dt$$

and

$$d(\ln N_t)=\frac{dN_t}{N_t}-\frac{c^2}{2}dt$$

Since

$$\frac{dN_t}{N_t}=\kappa dt+cdB_t$$

we see that

$$d(\ln N_t)=\frac{dN_t}{N_t}-\frac{c^2}{2}dt$$

and so

$$d(\ln N_t)=\kappa dt+cdB_t-\frac{c^2}{2}dt=(\kappa-\frac{c^2}{2})dt+cdB_t$$

The anti-derivative of $d(\ln N_t)$ can be written as $\ln\frac{N_t}{N_0}$:

$$\ln\frac{N_t}{N_0}=(\kappa-\frac{c^2}{2})t+cB_t$$

and finally:

$$N_t=N_0\exp((\kappa-\frac{1}{2}c^2)t+cB_t)$$
